# Irregular light schedules induce alterations on daily rhythms and gene expression in mice

**DOI:** 10.1101/2024.09.30.615784

**Authors:** Paula Berbegal-Sáez, Ines Gallego-Landin, Javier Macía, Olga Valverde

## Abstract

Synchronization of internal biological rhythms with external light-dark cycles is crucial for proper function and survival of the organisms, however modern life often imposes irregular light exposure, disrupting these internal clocks. This study investigated the effects of short-term shifted light-dark cycles on mice rhythmicity, and whether these alterations trigger molecular or behavioral changes. We evaluated locomotor activity, different behavioral domains and gene expression in the hypothalamus and medial prefrontal cortex. Despite non prominent behavioral impairments, such as anxiety or cognitive deficits, we observed a notable simplification in the locomotor activity patterns of the mice subjected to disrupted light-dark cycles. Molecular alterations included dysregulations in oscillations of core clock genes (*Cry2*, *Per2*) and disruptions in expression of genes involved in neuroplasticity, motivation, and stress responses, including *GluA1*, *Crhr2*, and *Vip* in both studied brain areas. Our study reveals that even brief light cycle shifts can disrupt circadian regulation at the molecular level, despite minimal behavioral changes. This molecular-behavioral discrepancy may suggest a complex adaptive response to drastic short-term light perturbations. Understanding the complex interplay between external light cues and internal biological rhythms regulation is crucial for mitigating the negative consequences of irregular light exposure on physiological processes and overall well-being.

## INTRODUCTION

The rhythmic cycle of light and darkness in nature is fundamental for synchronizing the internal clocks of living organisms. This intricate relationship between external light-dark (LD) cycles and internal biological rhythms is essential for regulating physiological processes and maintaining health and well-being. Light acts as one of the main synchronizing cues or zeitgebers that help regulate circadian clocks^1,2^. Thus, changes in the LD cycle scheme can significantly influence these internal rhythms, ultimately impacting an organism’s mood, cognition, and behavior^3,4^.

Circadian rhythms are orchestrated by the suprachiasmatic nucleus (SCN) in the hypothalamus, which acts as the body’s master pacemaker. It synchronizes internal clocks with environmental cues to regulate most biological functions such as sleep-wake cycles, hormone secretion, metabolism, and body temperature^5,6^. The core architecture of the mammalian circadian clock relies on a complex molecular network present in all cells of the body. Involving BMAL1 protein, and CLOCK protein, which together form the complex CLOCK:BMAL1. This complex heterodimer acts as a transcription factor promoting the expression of *Period* and *Cryptochrome* gene families. The resulting PER:CRY complex inhibits CLOCK:BMAL1, establishing a regulatory loop. Additional genes such as *Rev-erb* and *Rora* regulate BMAL1 expression in a secondary loop^7,8^. This intricate system manages the expression of various clock-controlled genes in a tissue specific manner^9^, influencing several physiological processes. Consequently, the de-synchronization of internal rhythms can lead to disturbances in numerous biological functions.

The disruption of natural LD cycles has garnered significant attention due to its profound effects on physiological functions and homeostasis. Understanding how alterations in light exposure can influence neurobiological processes is of crucial relevance. In this context, cognitive functions such as memory, attention, and decision-making are also intricately linked to the circadian rhythm. Studies have demonstrated that disruptions in the LD cycle can thereby impair cognitive performance by affecting neuronal activity and synaptic plasticity^10^. As a result, individuals exposed to irregular LD schedules, such as shift-workers, may experience a dysregulation of their internal biological rhythms which can ultimately result in different pathological conditions. Indeed, shift work, which disrupts the body’s natural circadian rhythm^11^, poses significant health risks, including accidents, and health issues such as obesity, cardiovascular disease, and mood disturbances^12,13^.

Typically, studies on photic entrainment patterns focus on LD cycles involving prolonged periods of light and darkness^14,15^. However, most organisms, including humans, do not experience continuous light exposure throughout the day. Providing insight into synchronization mechanisms in more translational settings is of utmost importance, since daily life has conditioned humans to be exposed to different hours of light throughout the day, often in irregular patterns that are not sustained over time. For instance, clinical studies have demonstrated that ten days of disturbed light conditions is enough to promote misalignment and alterations at a metabolic level potentially inducing a prediabetic state^16^. Additionally, research has linked exposure to light at night with an increased risk of mood disorders such as major depressive disorder, bipolar disorder or post-traumatic stress disorder severity^17–19^.

Within the central nervous system (CNS), misalignment of the molecular clocks is associated with dysfunction of several systems vital for maintaining homeostatic control^20^. Current studies have focused on relating pathological symptomatology of circadian disruption to specific neurobiological systems involved in affected behaviors. While it is understood that various neurotransmission systems, such as GABAergic^21^, glutamatergic^22^, and endocannabinoid^23^ signaling pathways, are involved in this disruption, the exact molecular interactions remain poorly understood, highlighting the need for further research. Particularly, systems related to stress regulation, metabolism, and motivated behaviors have shown sensitivity to circadian misalignment, ultimately influencing different stages of behavior^24–26^. These include the dopaminergic and opioid systems, alongside neuropeptides like AVP, VIP, NPY, orexins, and oxytocin^27–29^ all of which are involved in processes like energy balance, mood regulation, and social behavior^30^. Whether irregular light patterns can impact these systems is essential for understanding if such disruptions may contribute to pathological states.

This study aimed to investigate whether short-term irregular alterations in the LD cycle can disturb the internal clock and lead to behavioral impairments in mice. Specifically, it explores the relationship between aberrant LD cycles and changes in locomotor activity, mood, and cognition. We evaluated the circadian oscillations of the core clock genes as well as the expression of other genes involved in motivation, neuroplasticity, responses to stress, and metabolic control. Gene expression was investigated in the hypothalamus (HT), and medial prefrontal cortex (mPFC) a region crucial for cognitive tasks, decision-making and reward-related behaviors^31^.

Understanding these complex interactions is essential for developing effective strategies to prevent or mitigate the adverse consequences of irregular light exposure on physiological processes and to avoid pathological conditions resulting from disrupted biological clocks’ function.

## METHODS

### Animals

Male C57BL/6 mice aged 8 weeks were purchased from Charles Rivers (Lyon, France). The animals were housed in a controlled environment with *ad libitum* access to food and water, maintained at a temperature of 21 ± 1 °C, humidity levels of 55 ± 10%, and subjected to a 12:12 h LD cycle with lights off at 07:30 h. Prior to the beginning of any experimental procedures, mice underwent a minimum acclimatization period of one week to adapt to the new environmental conditions. All behavioral experiments were conducted when mice reached 10 weeks of age and during the dark phase under subdued red-light conditions.

The experimental protocols adhered strictly to the ARRIVE guidelines and regulations outlined by the European Community Council 2010/63/EU for animal experimentation and received approval from the local ethical committee (CEEA-PRBB).

### Experimental design

A total number of 134 male mice were distributed in three separate cohorts of animals used for distinct experimental procedures (Fig. 1). For all the studies, mice were divided into two groups: the control group, housed under a standard inverted LD cycle (dark phase from 7:30h – 19:30h; light phase from 19:30h – 7:30h), and the experimental group, subjected to a disrupted LD cycle (DLD) for 10 consecutive days. The distribution of animals for each study was as follows:

1. A first cohort of animals (n=5-6 mice/group) was used to assess daily locomotor activity patterns via continuous monitoring over a 48-hour period, beginning 24 hours post-DLD protocol. Following the monitoring period, the mice were subjected to molecular analyses.
2. A second cohort was utilized and divided into three subgroups to conduct different behavioral tests. A group of these animals underwent the elevated plus maze (EPM), saccharin preference test and splash test (n = 13 mice/group). A second group of mice was used to perform the social interaction test (n = 10 mice/group), forced swimming test (FST) and tail suspension test (TST) (n = 16 mice/group). A third group of mice was subjected to the location reference test and novel object recognition task (NOR) (n = 12-14 mice/group).
3. Lastly, a third cohort of animals (n= 3 mice/group and time point) was employed to obtain brain samples from the HT and mPFC to subsequent molecular analysis. Animals from the locomotor activity procedures were sacrificed to complete this batch (n= 2 mice/group and time point).

**Fig. 1.**
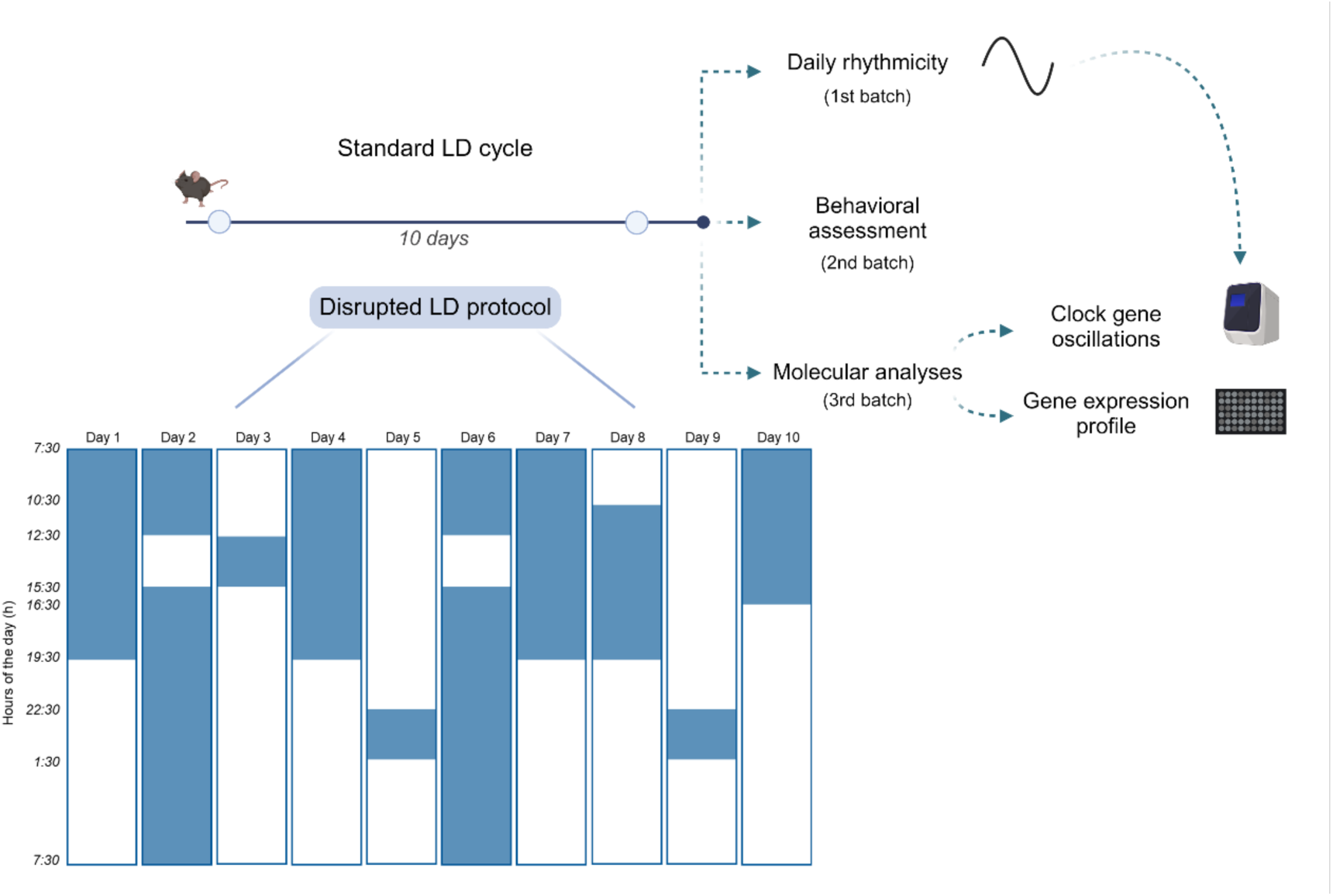
Representation of the experimental design with graphic schedule of disrupted light-dark protocol implemented. Mice were divided in two groups. The experimental group (DLD) underwent 10 consecutive days of disrupted light-dark protocol. Each column of the schedule represents one day of the protocol under which DLD mice were subjected. White regions inside each column represent the light hours per day and colored regions represent the hours of darkness. Hours of light changes are specified. Control mice were maintained under a standard inverted light-dark (LD) cycle (12L:12D). After the 10 days, animals were used for assessing rather daily rhythmicity, behavioral alterations, clock-gene oscillations, or molecular gene expression profile.

For the evaluation of daily oscillations in molecular expression, mice were euthanized every 4 hours over a 24-hour period. This approach allowed for a comprehensive evaluation of the temporal expression patterns of the main clock genes (*Bmal1, Clock, Cry2, Per2, Rev-erba*), providing insights into the circadian regulation of gene expression in the HT and mPFC. Also, an in-depth molecular profile was assessed in HT and mPFC using OpenArray^TM^ technology. This analysis evaluated a range of genes, including core clock genes, neurotransmission-related genes, and those involved in cognition, synaptic plasticity, motivation, stress responses, and metabolic processes (see Supplementary Table S2 for the complete gene list).

### Disrupted light-dark (DLD) cycle protocol

This protocol was designed to expose the animals to unpredicted LD cycles during 10 consecutive days. Based on a previous procedure^32^ with modifications, the DLD group experienced changes in the LD periods (Fig. 1). Over the course of 10 days, the experimental protocol implemented a variety of LD cycles to investigate their effects: standard inverted cycle of 12 hours of light followed by 12 hours of darkness on day 1 and day 4. Then some variations were introduced; on day 2 and day 6, there were 3 hours of light followed by 21 hours of darkness. On day 3, day 5 and day 9, the light period extended to 21 hours with only 3 hours of darkness. Finally, day 8 and day 10 featured 15 hours of light and 9 hours of darkness. After these 10 days, the animals returned to their standard inverted.

### Behavioral assessment

#### Actimetry

To evaluate spontaneous locomotion patterns throughout the LD cycle (12L:12D), each mouse was individually housed in a standard cage measuring 32 × 17 × 14 cm. Over a period of two consecutive days, the locomotor activity of each animal was monitored using the Panlab Infrared (IR) Actimeter (LE881 IR, Panlab s.l.u., Barcelona, Spain) and SEDACOM software, under 12L:12D cycle where ZT0 corresponded to the light phase onset (7:30 a.m). The motor activity (MA) of each mouse was quantified by calculating the number of counts per hour (MA/h). The time-series data obtained was analyzed using both, Kronos software and a novel mathematical approach based on Variation of Accumulated Entropy (VAE)^33^.

#### Reference Memory Test

Spatial reference memory was evaluated using a black Y-maze with two arms separated by 120° angles^34^. The experiment comprised training and testing sessions with a 1-hour inter-trial interval to assess short-term spatial memory. During training, one arm was closed off and labelled as the novel arm. In the test session, the mouse explored all three arms freely for 5 minutes. Visual cues were provided on each arm’s furthest wall to aid spatial recognition. The Smart software (Panlab s.l.u., Barcelona, Spain) recorded the time spent in each arm, and the preference ratio was calculated as: time spent in novel arm / total exploration time.

#### Novel Object Recognition task (NOR)

The Novel Object Recognition task, was conducted following the procedure outlined by previous studies from the group^35^. Black open boxes (24 cm × 26 cm × 15 cm) and plastic toys of similar size to mice were utilized. The task comprised three phases: habituation, training, and test sessions, all conducted under dim light conditions (15 lux). *Habituation:* Mice were acclimatized individually to the box for 10 minutes without objects. *Training Session:* Two identical objects (familiar objects) were introduced, and mice explored them for 10 minutes. *Test Session:* Three hours post-training, one familiar object was replaced with a novel one, and mice were allowed to explore for 10 minutes. The discrimination index was calculated as: (time spent exploring novel object - time spent exploring familiar object) / (time spent exploring novel object + time spent exploring familiar object) × 100.

#### Elevated Plus Maze (EPM)

This test utilized a black maze in a cross shape with two enclosed arms and two open arms, elevated 30 cm above the ground, following Portero-Tresserra et al., (2018)^36^. Mice were placed in the center of the maze and allowed to freely explore for 5 minutes. The SMART software (Panlab s.l.u., Barcelona, Spain) recorded parameters such as time spent in the open arms (OA), entries into the OA, and total distance. Results were expressed as a percentage of time spent in the OA (% time OA = time OA * 100 / (time OA + time CA)).

#### Tail suspension (TST)

Each mouse was suspended 50 cm above a workbench by placing adhesive tape 1 cm from the tail tip. Immobility time was measured manually with a digital chronometer over a period of 6 minutes following previous reports^37^.

#### Forced Swimming Test (FST)

Mice were placed in a glass cylinder 21 cm high with 15 cm of water^38^. Immobility time, defined as floating with minimal movements to maintain balance, was manually recorded over a 5-minute period.

#### Saccharin Preference Test

Following a protocol with minor modifications^39^, mice were individually housed with no prior food or water restriction. and exposed to a saccharin solution (0.33% w/v) and tap water for 48 hours. Bottle positions were alternated every 24 hours to prevent side preferences. Consumption of water and saccharin solution was assessed by weighing the bottles after 24 and 48 hours. Saccharin preference was calculated using the ratio: (intake of saccharin (g) / [intake of saccharin (g) + water intake (g)]).

#### Splash test

The Splash test was conducted in the home cage to minimize stress from an unfamiliar environment^40^. Mice were sprayed on their dorsal coat twice with a 10% sucrose solution diluted in tap water (∼2 × 0.6 ml). Self-grooming behavior duration (licking, stroking, and scratching) was manually recorded for 5 minutes as an indicator of motivation and self-care behavior.

#### Social Interaction Test

To assess social interaction, a three-chamber maze was utilized following established protocols^39^. Prior to the experiment, mice underwent a 5-minute habituation period in the maze. The evaluation consisted of two parts: *Sociability Assessment:* An unfamiliar mouse (M) was placed in one lateral chamber inside a metal holder, while an empty one (E) was placed in the opposite lateral chamber. The experimental mice were allowed to explore for 10 minutes. The sociability index was calculated based on the time spent exploring the mouse (TM) and the empty box (TE). Sociability Index (SI) = (TM - TE) / (TM + TE). *Social Novelty Assessment:* A novel mouse (M’) was placed inside the empty holder in the subsequent trial. The social novelty index was calculated by comparing the time spent exploring the familiar mouse (TM) and the novel mouse (TM’). Social Novelty Index (SNI) = (TM’ - TM) / (TM + TM’).

### Tissue collection and rt-qPCR

Mice were first sacrificed by cervical dislocation. Brain samples from HT and mPFC were collected using a 1 mm brain matrix. After their dissection, samples were immediately placed in dry ice and stored at −80°C. Sacrifices took place at different time-points corresponding to zeitgeber times [ZTs] 2, 6, 10, 14, 18, and 22. Total mRNA was extracted using TRIzol reagent following the manufacturer’s protocol, with RNA precipitation using isopropanol. cDNA synthesis was performed through reverse transcription of total mRNA. cDNA samples were employed for both clock gene expression time-point analysis through quantitative PCR (qPCR), and differential gene expression analysis with OpenArray™ technology. For the qPCR, the cDNA templates were combined with specific primers (see Supplementary Table S1) and SYBR Green PCR Master Mix. Also, the expression levels of target genes in each sample were normalized to *Gapdh* gene and expressed relative to the control group at ZT2. The fold change in gene expression between DLD and control animals was calculated using the 2-ΔΔCt method.

### OpenArray^TM^ technology

This technology allows the analysis of gene expression levels of several genes for multiple samples simultaneously. For that, 2.5 μl of each cDNA sample, from ZT14 and ZT18 time-point extractions, was combined with 2.5 μl TaqMan OpenArray^TM^ Real-Time Master Mix (Thermo Fisher #4462159) and loaded into a single well of a 384-well plate. Similar to previous studies^33^ custom OpenArray^TM^ plates (see Supplementary Table S2) were then automatically loaded using the AccuFill System and run in QuantStudio 12K. Amplification of the sequence of interest was normalized to endogenous reference genes, specifically, the geometric mean of *Actb*, *Gapdh*, and *B2m*. Fold-change values were calculated using the ΔΔCt method. Data were analyzed with the ThermoFisher Connect^TM^ cloud web tool.

### Data and statistical analysis

#### Data statistics

Normality (D’Agostino-Pearson and Kolmogorov-Smirnov tests), heteroscedasticity and homoscedasticity were assessed for all data sets. Significance was set at *p* < 0.05. For single factor, two-group analyses involving parametric variables we employed unpaired two-tailed Student’s *t-*tests. When an experimental condition followed a within-subject design a two-way ANOVA with repeated measures was conducted. Graphics and statistics were assessed in Graphpad software 8.0.

#### Entropy Divergence Analysis

In our analysis, we introduced a novel method called Variation of Accumulated Entropy (VAE), previously described in our previous work^33^ to assess differences in the complexity of information between two time series. Initially, the Discrete Fourier Transform was applied to the time series to obtain the set of *N* basic frequency components (harmonics) with associated phases and amplitudes. Each harmonic contains part of the information corresponding to the original time series. Subsequently, a subset of the first M harmonics was selected and the Shannon information entropy (E_M_) associated to this subset M was calculated. This process was repeated for each possible subset of M harmonics, ranging from M=1 to M=N. The procedure was applied to time series corresponding to DLD mice, i.e. 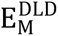 and to the CONTROL group, i.e. 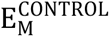. For each value M, *VAE(M)* was calculated as follows:

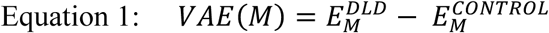

In Equation 1, *VAE(M)* quantifies the difference between both series in terms of informational complexity. If VAE values stabilize beyond a critical value *M*\*, it indicates that additional harmonics considered (i.e. *M*>*M*)* do not significantly provide additional information related to the intrinsic characteristics of the animals’ behavior. This stabilization occurs because these harmonics increase the informational complexity in both animal sets similarly, influencing the data series equally. This critical subset *M*\* contains essential information characterizing the activity patterns based on specific animal traits. In this context, VAE(M*) acts as a metric to quantify the differential complexity in internal rhythms between CONTROL and DLD animals. Applying the Inverse Discrete Fourier Transform to the subset composed of the first M* harmonics allows for the reconstruction of the time series without contributions unrelated to the intrinsic animal characteristics.

#### Kronos analysis

In our study, we employed Kronos, a computational tool developed by Bastiaanssen et al. (2023)^41^, and following previous studies methodology^33^ to analyze both locomotor activity patterns and biological gene expression rhythmicity. Kronos specializes in evaluating circadian rhythms within datasets, providing valuable insights into oscillatory responses. This software predicts sinusoid curves for variables based on specified periods, generating a kronosOut object with key outputs like variance proportion (average), p-values (significance for fit model), r-square (goodness-of-fit for sinusoidal data), acrophase, and amplitude. To compare rhythmicity between groups, Kronos utilizes a generalized linear model with user-defined predictors and interactions. It offers detailed model information and facilitates pairwise comparisons to distinguish overall differences and group-specific variations. Our analysis focused on essential 24-hour cycles crucial for health and physiological synchronization, revealing significant insights into oscillatory responses to internal and external factors.

## RESULTS

### Disruption of the LD cycle alters daily rhythms of locomotor activity in mice

Measurements of continuous locomotor activity for 2 days (Fig. 2a) revealed distinct behavioral patterns between the experimental groups. The oscillation analysis showed that activity peaks of the DLD mice were synchronized with the dark phase, thus presenting a daily oscillatory pattern (*p* < 0.01) similarly to control mice (*p* < 0.01) (Fig. 2b). However, these animals presented a slight advancement of the peak activity time (acrophase) respect to the control mice (control: 17.1 ZT; DLD: 14.5 ZT) and a minor increment of the amplitude (control: 146.1 a.u; DLD: 193.2 a.u). Eventually, statistical analysis using pairwise models indicated significant differences in rhythmic activity patterns between experimental conditions (*p* < 0.05).

**Fig. 2.**
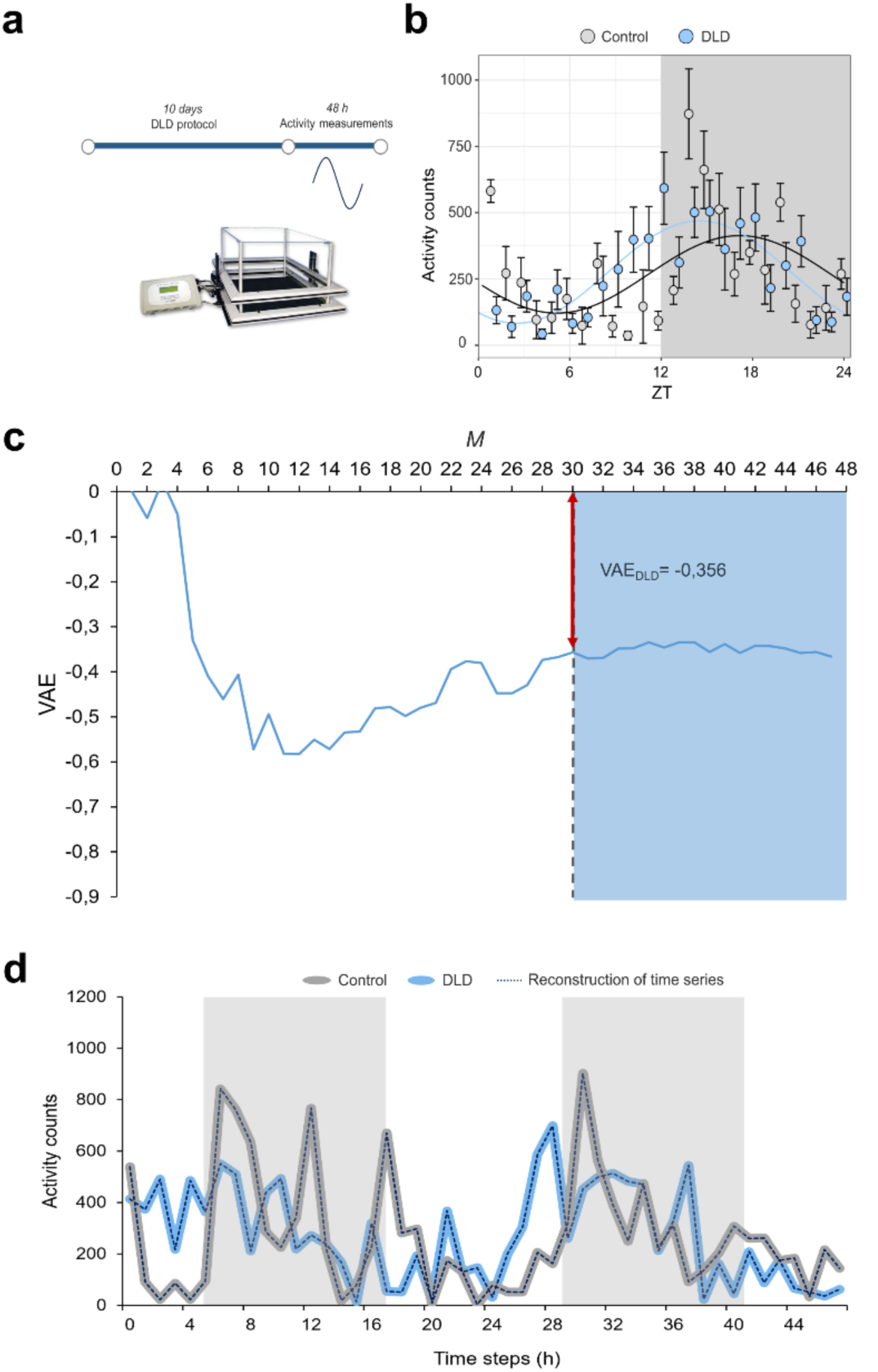
Assessment of daily arrhythmicity of disrupted light-cycle (DLD) mice. **(a)** Spontaneous locomotor activity in home cage recordings of two consecutive days under 12L:12D cycle**. (b)** Modelled sinusoidal curves of locomotor activity in control (grey) and DLD mice (blue). Grey area represents darkness hours of the day. Data is presented as mean + SD. **(c)** VAE values as a function of the number of Fourier components M used in their calculation. (**d)** Reconstruction of the time series using only the first M*=30 Fourier components, superposed to the mean of original locomotor activity data for control mice (grey line represent experimental data and dot-line represent the reconstruction series) and DLD mice (light blue line represent experimental data and dot-line represent the reconstruction series). Grey area represents dark hours of the cycle. (n = 5 mice per group).

To quantify these differences and assess changes in locomotor rhythmicity between experimental groups, we used the Variation of Accumulated Entropy (VAE) metric as a novel approach to quantify alterations in daily rhythms^33^. Through the decomposition and mathematical analysis of our experimental data, VAE was computed. Fig. 2c illustrates the VAE values as a function of the number of Fourier components (M). In the initial region (white area), VAE values exhibit significant fluctuations with increasing components. This variability suggests that each component of the DLD subset conveys unique information specific to either the control or DLD group, reflecting group-specific characteristics. In the subsequent region (colored area), starting from M*= 30, VAE values displayed minimal fluctuations, indicating that additional Fourier components provided similar information across both groups, suggesting shared factors rather than group-specific attributes. Here, VAE= −0.356 value at M*= 30 quantifies the difference in pattern complexity between the two groups. A negative value reflects that control mice exhibit more intricate daily dynamics compared to DLD mice. Figure 2d depict the original time series Control(t) and DLD(t) along with the reconstruction of the time series using only the first M*= 30 Fourier components. In both groups, this reconstruction closely aligns with the original series, indicating that these 30 components capture the primary features of the locomotor activity patterns.

### Molecular clock machinery is altered within the HT and mPFC of DLD mice

The use of Kronos software for gene oscillation analysis allows for a comprehensive evaluation of rhythmicity, revealing distinct gene oscillation patterns between the control and DLD groups as reported regarding different parameters presented in Table 1. Figure 3 illustrates regression curves for gene expression data over time in both groups and for *Bmal1*, *Clock*, *Per2*, *Cry2*, and *Rev-erba* clock genes.

**Fig. 3.**
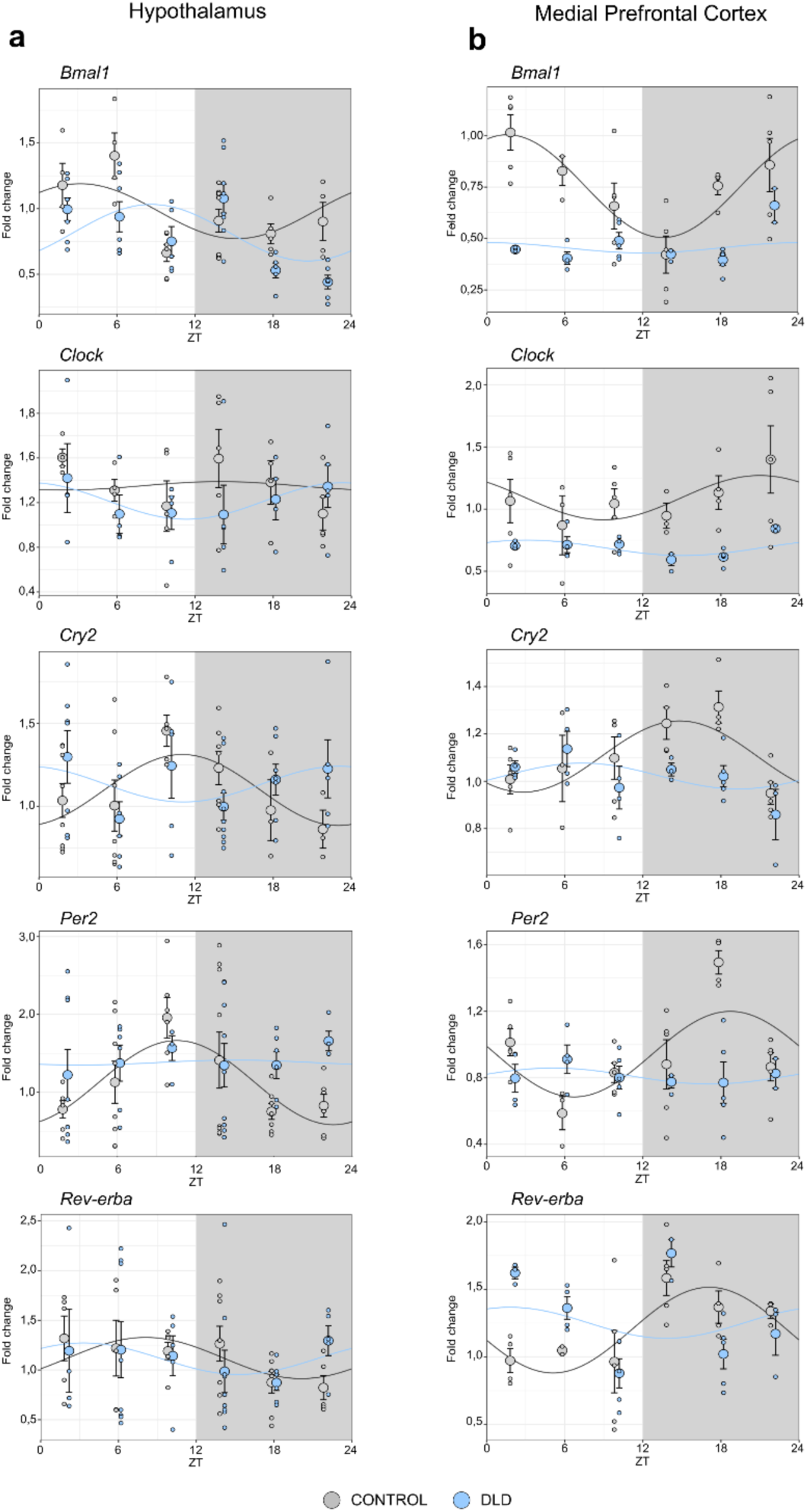
Clock gene expression desynchrony and arrhythmicity in the HT and mPFC of DLD mice. Expression levels (relative to ZT 2 control group) of clock genes - *Bmal1, Clock, Cry2, Per2, Rev-erbα* – in different ZT time points (n= 4-5 mice per group and time point), within the **(a)** Hypothalamus and **(b)** Medial Prefrontal Cortex. Kronos software returned the graphs of the cosine-fitted curves of control and DLD mice, and subsequent analysis. Period =24h, 12h light (white zone) 12h dark (grey zone). ZT, Zeitgeber time, where ZT0 lights turn on.

**Table 1.**
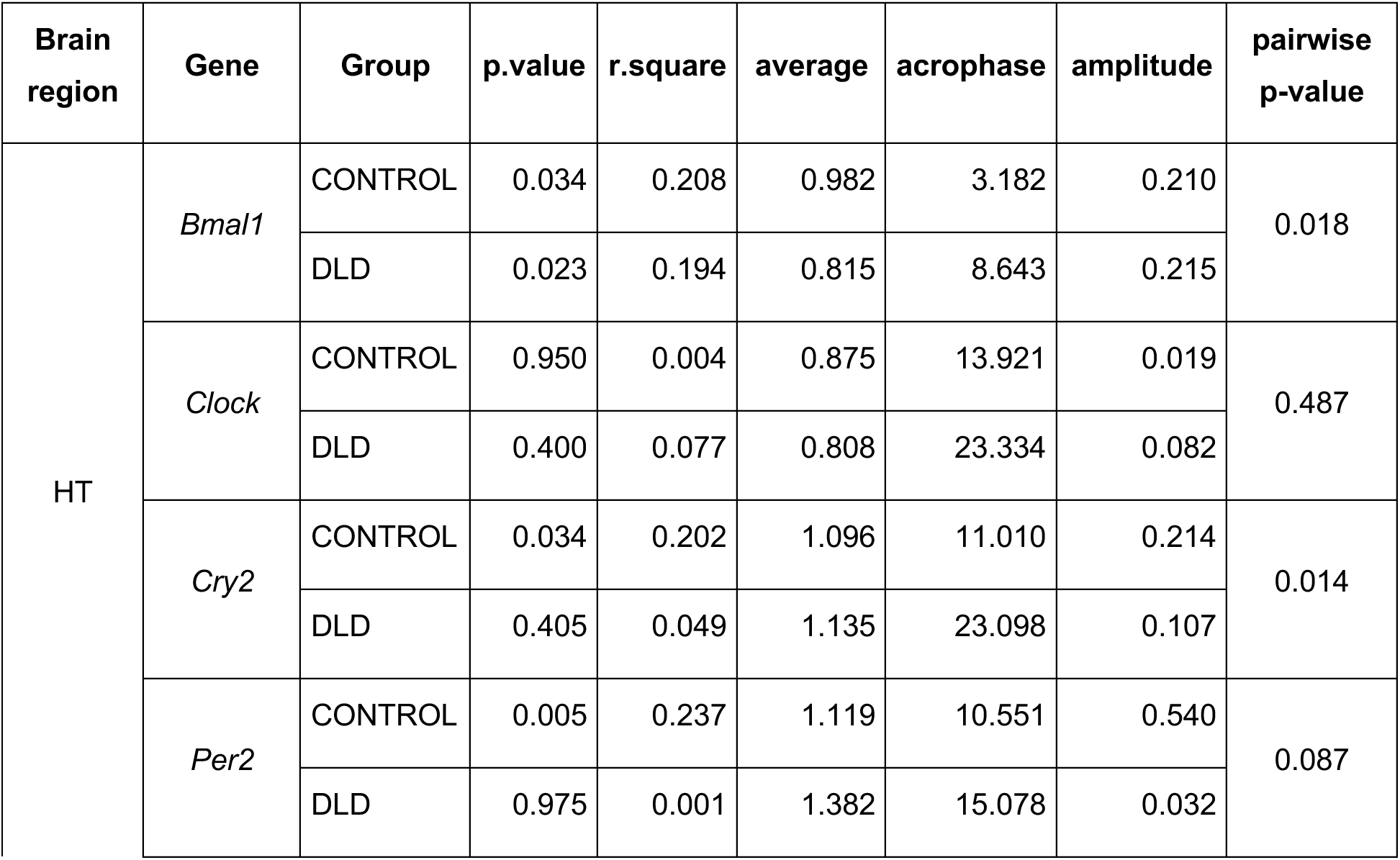
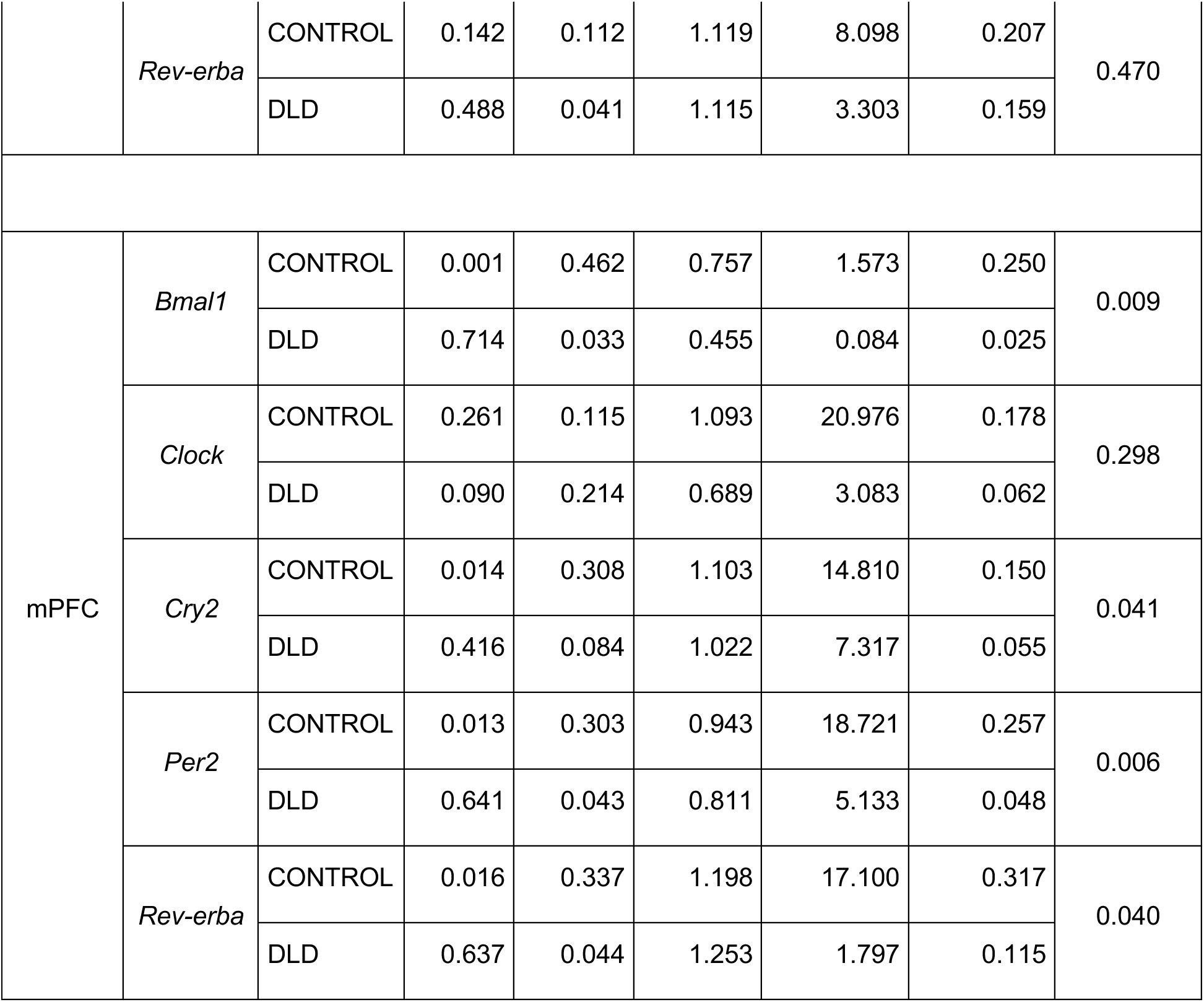
Circadian rhythmicity analysis output of clock genes.

Significant oscillations were observed in the HT (Fig. 3a) and the mPFC (Fig. 3b) for the control group for the genes *Bmal1*, *Cry2*, *Per2*, and only in the mPFC, for the *Rev-erba*. *Clock* did not exhibit oscillations in either the control group or the DLD group in either of the regions studied, according with its constitutive expression pattern^42,43^.

In the experimental group a loss of rhythmic oscillations was identified for *Cry2* and *Per2* in the HT, and for *Bmal1*, *Cry2*, *Per2* and *Rev-erba* in the mPFC. The pairwise comparisons (*p* < 0.05) indicated the significant difference of gene oscillation patterns of all these genes between both groups (Table 1). Regarding *Bmal1*, its oscillation was present in the HT for DLD mice. However, the expression pattern is reportedly different compared to the control group (pairwise *p* < 0.05). The analysis particularly reports a shift in the acrophase (Control: 3.182 ZT; DLD: 8.643 ZT).

Overall, the molecular machinery of the DLD animals was affected in both brain regions with special impact on the mPFC, in which the oscillation of the studied clock genes but *Clock*, was disrupted.

### Irregular disruption of LD cycle does not evidence significant behavioral alterations

Different behavioral domains were evaluated to investigate the impact of disrupted LD cycle on mice.

Regarding the cognitive domain, the reference memory test (Fig. 4a) (t25 = 0.152; *p* = 0.879), and NOR task (Fig. 4b) (t21 = 0.385; *p* = 0.704), showed comparable results between groups.

**Fig. 4.**
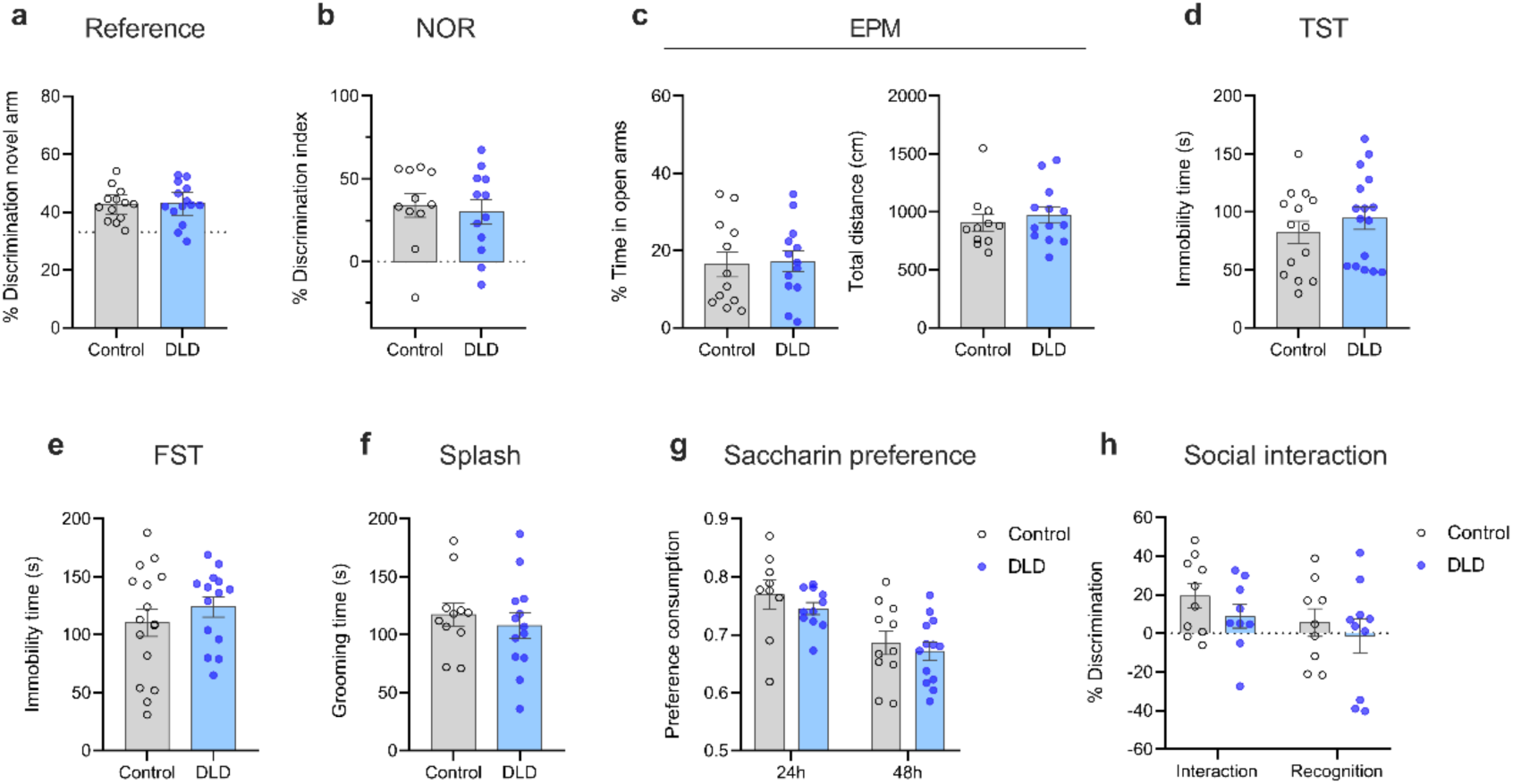
Behavioral assessment of the effects promoted by a disrupted light-dark cycle. Mice with disrupted light-dark (DLD) cycles exhibit no significant effects in **(a, b)** cognitive function measured through the Reference Memory Task and Novel Object Recognition (NOR), **(c)** anxiety levels as assessed by the Elevated Plus Maze (EPM), **(d, e)** despair-like behavior analyzed with the Tail Suspension Test (TST) and Forced Swim Test (FST), **(f, g)** anhedonia assessed through Saccharin preference and Splash tests and **(h)** sociability evaluated using the Social Interaction Test. Data are presented as mean ± SEM, with n = 10-16 mice per group. Statistical analysis was performed using unpaired t-tests or ANOVA.

Anxiety levels assessed by the EPM (Fig. 4c) showed no significant differences between the control group and the DLD group in terms of percentage of time spent in open arms (t23 = 0.204; *p* = 0.839) and total distance (t22 = 0.6216; *p* = 0.540). Similarly, assessing despair-like behavior, no significant difference in immobility time was observed between control and DLD group in neither the TST (Fig. 4d) (t28 = 0.871; *p* = 0.391) nor in the FST (Fig. 4e) (t28 = 0.896; *p* = 0.377).

Regarding anhedonia-like state, the splash test revealed no significant differences in grooming time (t22 = 0.619; *p* = 0.541) (Fig. 4f). Furthermore, the saccharin preference test (Fig. 4g), demonstrated no significant variation in saccharin preference consumption in *Group* effect (F(1, 23) = 1.063; *p* = 0.3133), whereas *Days* effect (F(1, 17) = 24.53; *p* < 0.0001) showed significant differences, suggesting increased saccharin consumption throughout the test in both groups.

Finally, in the same line, analysis of the social interaction test data (Fig. 4h) showed no significant differences for neither social interaction (t17 = 1.179; *p* = 0.254) nor social recognition (t17 = 0.620; *p* = 0.543) between DLD and control groups.

### Disrupted LD cycle alters the expression of core clock genes and related systems in HT and mPFC

Genes showing an absolute change greater than 1.3-fold in either direction with a p-value smaller than 0.05 exhibited differential expression in the DLD group (Fig. 5). Concrete values for significantly different expressed genes, of both fold change and p values, are presented in Fig. 5 and Supplementary Table S2.

**Fig. 5.**
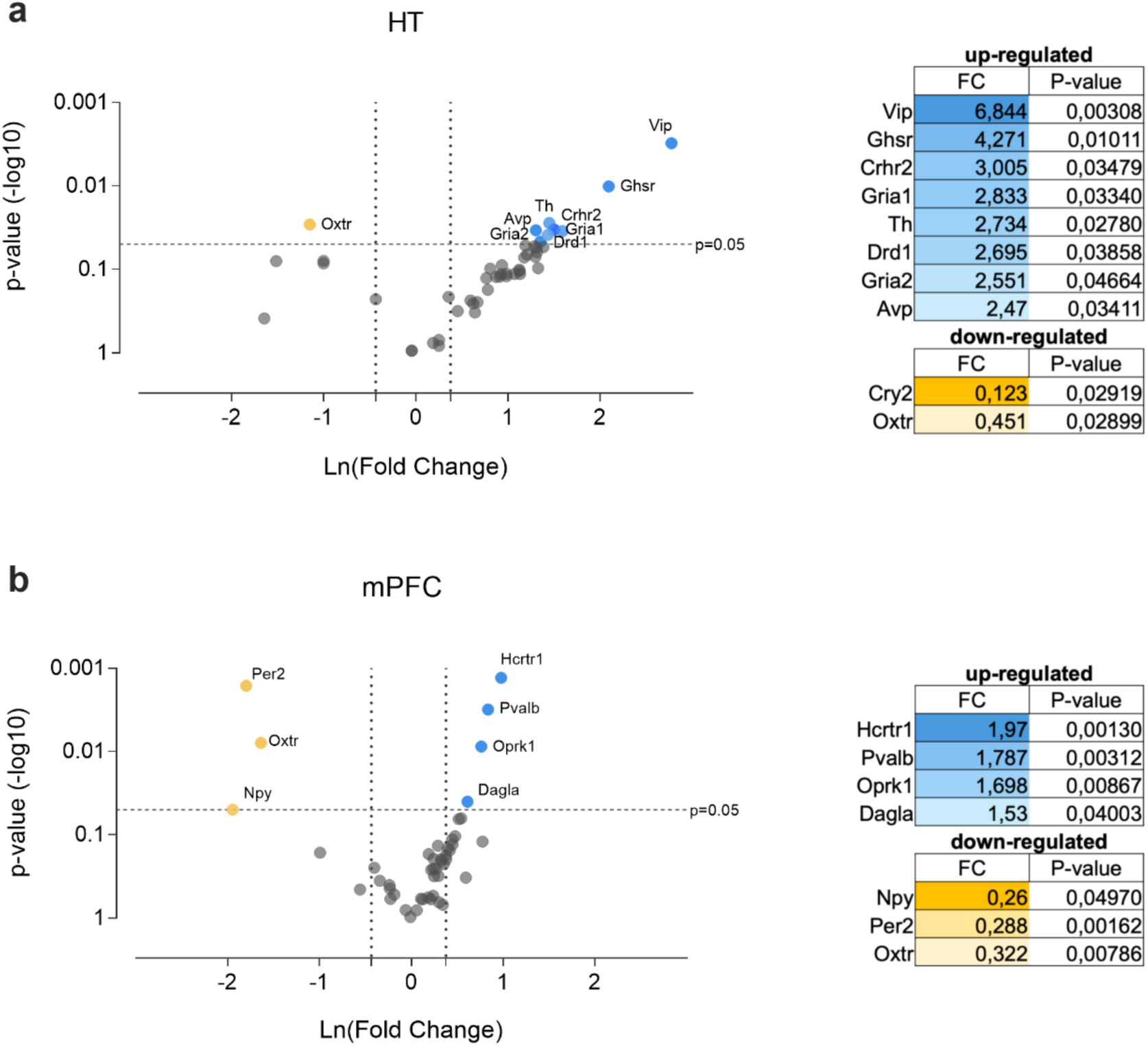
Differentially expressed genes of DLD group compared to non-disturbed light-cycle mice. Volcano plots represent the differential expression of 52 genes between the disrupted animals DLD (n = 8) and control group (n = 8), assessed in OpenArray^TM^ analysis of mRNA levels within the **a.** HT and **b.** mPFC. The graphs illustrated the association between fold change and significance in two groups, using a student’s t-test. The y-axis showcases the negative log10 of p-values from the t-tests (where the horizontal slider corresponds to a p-value of 0.05), while the x-axis represents the difference in expression between the experimental groups as Ln fold changes. Vertical sliders indicate mRNAs as either up-regulated (located in the right area, denoted by blue dots) or down-regulated (situated in the left area, represented by yellow dots) from a fold change (FC) of 1.3. The genes that displayed both an absolute change greater than FC1.3 in either direction, and a *p*-value <0.05 were identified as the differentially expressed genes.

Clock genes were affected in both regions, with *Cry2* downregulated in the HT and *Per2* downregulated in the mPFC.

Additionally, in HT (Fig. 5a), we reported a dysregulation of multiple genes related to different systems. Thus, we observed a downregulation of the oxytocin receptor (*Oxtr)*, while some neuropeptides (*Vip, Avp*), genes involved in metabolic control, as ghrelin receptor (*Ghsr*), and genes involved in stress responses, as corticotropin-releasing hormone receptor 2 (*Crhr2*), were upregulated. Eventually, glutamate receptors *Gria1* and *Gria2* along with genes related to dopamine system such as dopamine receptor 1 (*Drd1)* and tyrosine hydroxylase (*Th*), were also upregulated in DLD animals compared to control.

Within the mPFC (Fig. 5b), we identified dysregulation in genes involved in different systems. In this area we also find a downregulation of *Oxtr* and the neuropeptide Y (*Npy)*. The hypocretin receptor 1 (*Hcrtr1*) from orexinergic system, parvalbumin (*Pvalb*) involving GABAergic transmission, the opioid kappa receptor (*Oprk1*) and diacylglycerol lipase (*Dagla*) were found to be upregulated in DLD mice.

These findings highlight a widespread dysregulation of clock genes and related neurobiological systems in the DLD group across different brain regions.

## DISCUSSION

The present study aimed to investigate whether acute, irregular, and unpredictable shifts in the LD cycle disrupts the internal clock and leads to behavioral impairments in mice. Our findings revealed that animals exposed to short term disruption of LD cycles showed changes in their daily rhythms and disturbances in clock gene oscillations within the HT and mPFC. While behavioral alterations were not prominent, DLD condition triggered substantial molecular changes including variations in some neuropeptide’s expression, clock genes, and genes involved in neurotransmission pathways, as well as signaling pathways related to motivation, neuroplasticity, stress responses and metabolic regulation.

Locomotor activity analyses revealed that DLD mice experienced an advancement in the onset of their activity phase, *i.e.* acrophase, and showed different activity oscillation patterns compared to control mice. This suggests that the aberrant LD cycle affects the circadian machinery, shifting the peak of locomotor activity, and ultimately causing misalignment with external cues. The employment of VAE analysis allowed us to quantify differences of activity patterns providing additional information that enhances the consistency in quantifying circadian dysregulations compared to previous procedures^44^. The VAE = −0.356 highlights the distinction between both groups, indicating a simplification of the activity pattern compared to control group. If the locomotor patterns in DLD mice were similar to the control group, the VAE would approach zero. This finding implies that the shifted light protocol led to a reduction in the complexity of the animals’ activity cycles, potentially indicating a disruption in the circadian organization. This simplification might suggest a reduced ability of the organisms to adapt and maintain circadian rhythms under this particular environmental perturbations. Previous studies utilizing VAE analysis applied to more impactful conditions, such as the deletion of BMAL1, resulted in a more complex mice’ activity pattern^33^, suggesting that different types of circadian disruption may result in alterations od daily patterns rather increasing or reducing their complexity. While the underlying mechanisms of the simplification or complexity of the pattern remain unclear, these findings raise questions regarding the causes of internal rhythm disruption and underscores the value of continued research on further properties of circadian rhythms such as the complexity of the activity pattern. Several different protocols of aberrant LD cycles demonstrated to induce changes in locomotor activity often accompanied by alterations in behavioral domains. Clark et al. (2020)^32^ found a disruption in activity after acute shifts in the LD cycles, which promoted fear extinction impairments. Similarly, protocols mimicking jet lag conditions were associated to disruptions in activity phases and were found to negatively impact spatial and recognition memory^45^. Together these findings highlight the broad impact of light cycle perturbations on behavior.

However, in contrast to the locomotor activity results, we did not observe significant signs of behavioral disruptions in our experimental group across various domains, including recognition and spatial memory, anxiety-like behavior, sociability, and motivation. This suggests that short-term disruptions in the LD cycle are insufficient to significantly affect behavior beyond locomotor activity. In literature, the most reported affected behavior by irregular lightning conditions is fear memory extinction^32,46^, a parameter that the present work does not explore. Nonetheless, other LD schemes have yielded results similar to ours, where mice showed changes in activity but not in other behavioral domains^47,48^. This discrepancy in findings suggests the existence of adaptive mechanisms that may confer resilience to some environmental perturbations, underscoring the organism’s ability to adapt to shifts in the LD cycle for proper functioning and homeostasis. In fact, some research in humans suggests that fast-rotating shift schedules are generally more tolerable for shift workers compared to slower-rotating schedules^46^, likely because the circadian machinery adaptatively does not entrain anymore with external light cues, but follow its internal rhythms when rapid changes significantly vary from naturally occurring LD cycles. Organisms may perceive very irregular light conditions as inefficient for adaptation and instead prioritize internal cues. Nonetheless, it is possible that chronic circadian disruption or more severe or prolonged LD changes, may eventually manifest in more significantly behavioral changes^49^, such as hippocampal learning and mood disregulation^50^. While our investigation focused on acute circadian disruption, the potential long-term consequences of chronic misalignment need further examination. Such research could provide crucial evidence supporting the link between persistent circadian misalignment and the onset of disease.

Despite our findings suggesting resilience to behavioral disruptions under short-term DLD exposure, the observed molecular alterations might render the organism more susceptible to other insults or stressors, this is, potentially eliciting behavioral changes that have not yet been observed^51,52^. Subsequent studies could introduce environmental or physiological challenges after the DLD protocol to reveal potential vulnerabilities and offer deeper insights into how circadian misalignment impacts behavior and cognitive functions under more challenging conditions. Nevertheless, the current findings highlight the importance of considering both molecular and behavioral outcomes in circadian rhythm research, as molecular changes may precede observable behavioral alterations. This is particularly significant, prompting us to further investigate potential molecular alterations that hat could contribute to broader system dysregulations.

In our study, we found changes in clock gene oscillations in the DLD group, specifically the loss of rhythmicity of both *Cry2* and *Per2* within the HT. As core molecules in cellular timekeeping, their altered oscillation patterns indicate disruptions in the circadian machinery. We observed these alterations despite the appropriate oscillation of *Bmal1*, a direct controller of their expression. The differences in patterns, likely due to a shift in the acrophase, could contribute to disruptions in the circadian system and, therefore, to desynchrony among tissues. Comparable changes in core clock genes in the hypothalamus have been observed in mice exposed to mild changes in environmental lighting conditions, such as dim light at night^53^. While that study found circadian patterns in Clock expression, our observations of *Clock* as a constitutively expressed gene align with previous reports^42,43^, which supports the consistency of our findings.

Furthermore, we report disruptions in the oscillations of core clock genes *Bmal1, Per2, Cry2* and *Rev-erba* in the mPFC, revealing daily rhythmic disruptions in this region. This prompts the question of whether similar disturbances may occur in other brain areas, as previously seen in hippocampus, where comparable irregular light conditions have been shown to affect clock gene expression^26^. This holds significant importance, since alterations in non-SCN cellular clocks are more susceptible to genetic and environmental influences^49^ compared to the SCN network clock. Said susceptibility may result in impairments across various processes beyond maintaining internal synchrony. Our results mirror the flattened clock gene expression observed in mPFC after irregular light cycles^26^ and, interestingly, align with previous studies reporting a loss of rhythmicity—rather than a phase shift—under jet-lag-like schedules^54^. Possibly, the variability of light changes in our protocol may have contributed to the inability of internal clocks to properly synchronize with any consistent oscillating pattern.

To understand whether experimental mice experienced further molecular alterations linked to internal rhythm changes, we conducted a deeper molecular analysis of the HT and mPFC. The distinct gene expression profile observed in DLD mice, indicated the affectation of various physiological systems, not only of different clock genes, but also neuropeptide expression, and genes related to neuroplasticity, motivation, and stress-related responses. This molecular changes may also reflect the internal misalignment and disruption, considering that nearly 20% of mammalian genes are regulated by circadian machinery^55^.

Among the affected hypothalamic systems, we report an up-regulation of genes involved in the glutamatergic activity such as *Gria1* and *Gria2* genes, which encode subunits of glutamate AMPA receptors. These receptors are important for neuronal excitability and neuroplasticity and their dysregulation can disrupt downstream signaling pathways related to motivation^20^. Disruptions of GluA1 subunit (*Gria1* gene) have been linked to circadian disruption and aberrant responses to environmental cues^56^. To our knowledge, this is the first report of glutamate receptor subunit changes induced by alterations in light schedules, confirming the influence of environmental light alterations in glutamatergic regulation. Additionally, the dopaminergic system, crucial in motivational responses, was affected in DLD mice through changes in *Th* and *Drd1* gene expression. These genes are regulated in a circadian manner^57,58^, further supporting our findings on rhythmic dysregulation in organisms subjected to an aberrant LD cycle. Even metabolic control appears to be affected by circadian disruption^25^, consistent with the reported overexpression of *Ghsr* in the DLD mice. Furthermore, *Crhr2* involved in stress regulation through the activation of the hypothalamic-pituitary-adrenal (HPA) axis, is also subjected to circadian rhythm control under physiological conditions^59^. Activation of corticoid receptors has been linked with the control of *Per* expression in both central nervous system and peripheral tissues^60^. The reported different *Per2* oscillations in DLD mice align with a *Crhr2* dysregulation and are consistent with other models of circadian disruption that show alterations in corticosteroid levels^24^, prompting the adrenal clock as a crucial player in the regulation of changes induced by aberrant light exposures^61^.

Further, altered expression of *Vip* and *Avp* within the HT suggests potential disruptions in SCN function, indicating possible dysregulation of the circadian system. These are the two primary neuropeptides that signalize the neurotransmission between the core and the shell of the SCN, and thus are fundamental in synchronizing the central circadian clock with external LD cycles^27^. Changes in these genes may trigger a cascading effect on the regulation of numerous functions, including metabolic processes, as well as stress control within the HT.

Moreover, dysregulation of signaling pathways extends to the mPFC. In this brain area other neuropeptides including *Npy,* which plays a crucial role in stress response and mood regulation, was downregulated. While high levels of NPY in the mPFC have been associated with phenotypic resilience, decreased expression, as observed in our study, are linked to mood impairments^62^. This suggests that reduced levels of *Npy* expression may be a risk factor for developing depressive-like states. Within this area, DLD exposure also promoted changes in genes involved in GABAergic transmission, notably the upregulation of *Pvalb*, previously associated with anxiety-like behaviors^63^. In addition, *Oprk1* and *Dagla*, which are involved in the activity of the opioid and the endocannabinoid systems respectively, were differentially expressed in DLD mice. These systems regulate signaling from the mPFC to other brain regions, and their dysregulation could potentially contribute to dysphoric states^64^ while impacting emotional and cognitive processing^65^. Furthermore, the oxytocin system was also affected, as indicated by changes in *Oxtr* expression. This system conveys light information to regulate feeding behavior and synchronize internal rhythms^29^, and also responds to the presence of stressful stimuli^66^. Overall, many systems are reportedly affected following environmental light changes. The observed alterations in gene expression likely result from the intrinsic dysregulation of daily rhythms in mice subjected to DLD cycles and suggest a state of physiological dysregulation that may increase vulnerability to future challenges.

Notably, these molecular alterations occurred despite the absence of observable behavioral changes, suggesting potential compensatory mechanisms or a threshold effect in the translation of gene expression changes to behavioral outcomes. This highlights the intricate interplay between circadian rhythms and various physiological processes as well as the importance of molecular analysis in understanding the subtle effects of circadian disruption, even in the absence of overt behavioral changes.

Recent research evidences significant sex differences in physiological responses, behavioral patterns, and immune function when circadian rhythms are disrupted^67^. For example, Walker et al. (2020)^68^ demonstrated that even brief exposure to dim light at night alters brain physiology and behavior differently in male and female mice. Furthermore, it is clear that the estrous cycle in females introduces an additional layer of complexity to circadian studies due to its influence on gene expression patterns^69^. This cyclical variation requires careful experimental design and analysis to account for hormonal fluctuations. Acknowledging such differences in the two populations, in the present study we have focused exclusively on male mice. Nonetheless, future research should aim to replicate these experiments on female mice, with a methodology specifically designed to address the intricacies of female physiology. This would provide a more comprehensive understanding of how irregular light schedules affect daily rhythms and gene expression across sexes. Additionally, such research could reveal potential sex-dependent vulnerabilities or resilience mechanisms to circadian disruptions.

In conclusion, this research debates the intricate interplay between environmental light cues, biological rhythms, and their molecular and behavioral outcomes. Our findings show that disrupted LD cycles can perturb the biological clock’s rhythmicity and alter the expression of genes crucial for neuroplasticity, motivation, and neurotransmission. The observed molecular changes, coupled with the absence of significant behavioral changes, underscore the adaptive capacity and resilience of biological systems. This research contributes to our understanding of circadian biology and raises important questions about the potential long-term consequences of exposure to irregular chronic circadian disruption.

## Data availability statement

Further information and requests for data, resources and reagents should be directed to and will be fulfilled by the lead contact, Dr. Olga Valverde upon request.

## Acknowledgements

This work was supported by a grant from the Ministerio de Ciencia e Innovación (PID2022-136962OB-100 - MCIN/AEI/10.13039/501100011033 and PID2020-119538RB-I00 -MCIN /AEI/10.13039/501100011033), ESF “A way of making Europe”, by Ministerio de Sanidad, Delegación del Gobierno para el Plan Nacional sobre Drogas (#2023/005) and Fondos de Recuperación, Transformación y Resiliencia (PRTR) Unión Europea (#Exp2022/008695), and by the Generalitat de Catalunya, AGAUR (#2021SGR00485). P.B-S is supported by FI-AGAUR grant from the Generalitat de Catalunya (#2021FI-B00205). I.G-L. is supported by a grant from the Ministerio de Ciencia e Innovación (#PRE2020-091923). O.V. is recipient of an ICREA Academia Award (Institució Catalana de Recerca i Estudis Avançats, Generalitat de Catalunya).

## Author contributions

P.B-S.: Conceptualization, Data curation, Formal analysis, Investigation, Methodology, Visualization, Writing - original draft, Writing - review & editing. I.G-L.: Data curation, Formal analysis, Investigation, Methodology, Writing - review & editing. J.M.: Conceptualization, Formal analysis, Methodology, Writing - review & editing. O.V.: Conceptualization, Methodology, Formal analysis, Funding acquisition, Supervision, Visualization, Writing - review & editing.

## Additional information

Authors declare no conflict of interests.

## Notes

### Competing Interest Statement

The authors have declared no competing interest.

